# FLAME: a web tool for functional and literature enrichment analysis of multiple gene lists

**DOI:** 10.1101/2021.06.02.446692

**Authors:** Foteini Thanati, Evangelos Karatzas, Fotis A. Baltoumas, Dimitrios J. Stravopodis, Aristides G. Eliopoulos, Georgios A. Pavlopoulos

## Abstract

Functional enrichment is a widely used method for interpreting experimental results by identifying classes of proteins/genes associated with certain biological functions, pathways, diseases or phenotypes. Despite the variety of existing tools, most of them can process a single list per time, thus making a more combinatorial analysis more complicated and prone to errors. In this article, we present FLAME, a web tool for combining multiple lists prior to enrichment analysis. Users can upload several lists of preference and use interactive UpSet plots, as an alternative to Venn diagrams, to handle unions or intersections among the given input files. Functional and literature enrichment along with gene conversions are offered by g:Profiler and aGOtool applications for 197 organisms. In its current version, FLAME can analyze genes/proteins for related articles, Gene Ontologies, pathways, annotations, regulatory motifs, domains, diseases, phenotypes while it can also generate protein-protein interactions derived from STRING. We have herein validated FLAME by interrogating gene expression data associated with the sensitivity of the distal part of the large intestine to experimental colitis-propelled colon cancer. The FLAME application comes with an interactive user-friendly interface which allows easy list manipulation and exploration, while results can be visualized as interactive and parameterizable heatmaps, barcharts, Manhattan plots, networks and tables.

**Availability:** FLAME application: http://flame.pavlopouloslab.info

**Code:** https://github.com/PavlopoulosLab/FLAME

## INTRODUCTION

Functional enrichment analysis is a method to identify classes of bioentities in which genes or proteins were found to be over-represented. This type of analysis can aid researchers to reveal biological insights from various -Omics-experiments and interpret gene lists of interest in a biologically meaningful way. For this purpose, several applications have been proposed (1, 2). Established tools include: g:Profiler (3), Panther (4), DAVID (5), WebGestalt (6), EnrichR (7), AmiGO (8), GeneSCF (9), AllEnricher (10), aGOtool (11), ClueGo (12), Metascape (13), NeVOmics (14), GSEA (15), GOrilla (16), Fuento (17) and NASQAR (18). Most of them are offered as web applications and among other functionalities, they associate overrepresented genes with GO terms (19), or pathways (20–23). Nevertheless, (a) they often differ in the organisms, the identifiers and the backend database they support, (b) they come with their own way of reporting results (mostly lists or static representations) and (c) they are often unable to handle and compare multiple lists for a more combinatorial analysis.

To address such critical issues, in this article, we present FLAME, a web application which allows the combination of multiple input gene lists, and their parallel exploration and analysis with the use of interactive UpSet plots. FLAME integrates g:Profiler and aGOtool for an accurate and always up-to-date functional and literature enrichment analysis, and also utilizes STRING’s API (24) to generate interactive protein-protein interaction (PPI) networks. A major advantage is that FLAME follows a visual analytics approach to allow users to adjust and parameterize the reported results with the engagement of heatmaps, barcharts, Manhattan plots, networks and tables, thus making information easier to be absorbed, comprehended and interpreted, and knowledge easier to be extracted and exploited.

## METHODS

### Input

FLAME allows the uploading of multiple gene lists as separate files, or as pasted text. In the online version, FLAME can accommodate up to ten active gene lists with a file size smaller than 1MB each, a limitation which can be bypassed by downloading the application from GitHub and running it locally after editing the value of the corresponding variable (FILE_LIMIT configuration variable in global.R and shiny.maxRequestSize option in ui.R). The input text can be imported in comma-, tab- or line-separated format. In this version, FLAME supports 197 organisms and several identifiers, such as gene IDs (proteins, transcripts, microarray IDs, etc), SNP IDs, chromosomal intervals and term IDs, in accordance with the gconvert function of the gprofiler2 library. Different lists are not allowed to get the same name with options for renaming and deleting them being suitably provided. Once uploaded, list contents can be seen in interactive searchable tables.

### List manipulation with the use of UpSet plots

After uploading the gene lists of interest and prior to analysis, UpSet plots can be generated to show possible intersections, unions and distinct elements between the selected lists. The UpSet plot is a sophisticated alternative of a Venn-diagram and is prefered for many sets (>5) where a Venn diagram becomes incomprehensive. In Figure 1, we demonstrate a simple example with three lists, each containing 100 random genes, and visualize the various UpSet plot options in relation to a Venn diagram (Figure 1, A-D). Similarly, in Figure 1E, we describe the distinct intersections among seven gene lists, a task which cannot be drawn as a Venn diagram.

**Figure 1.**
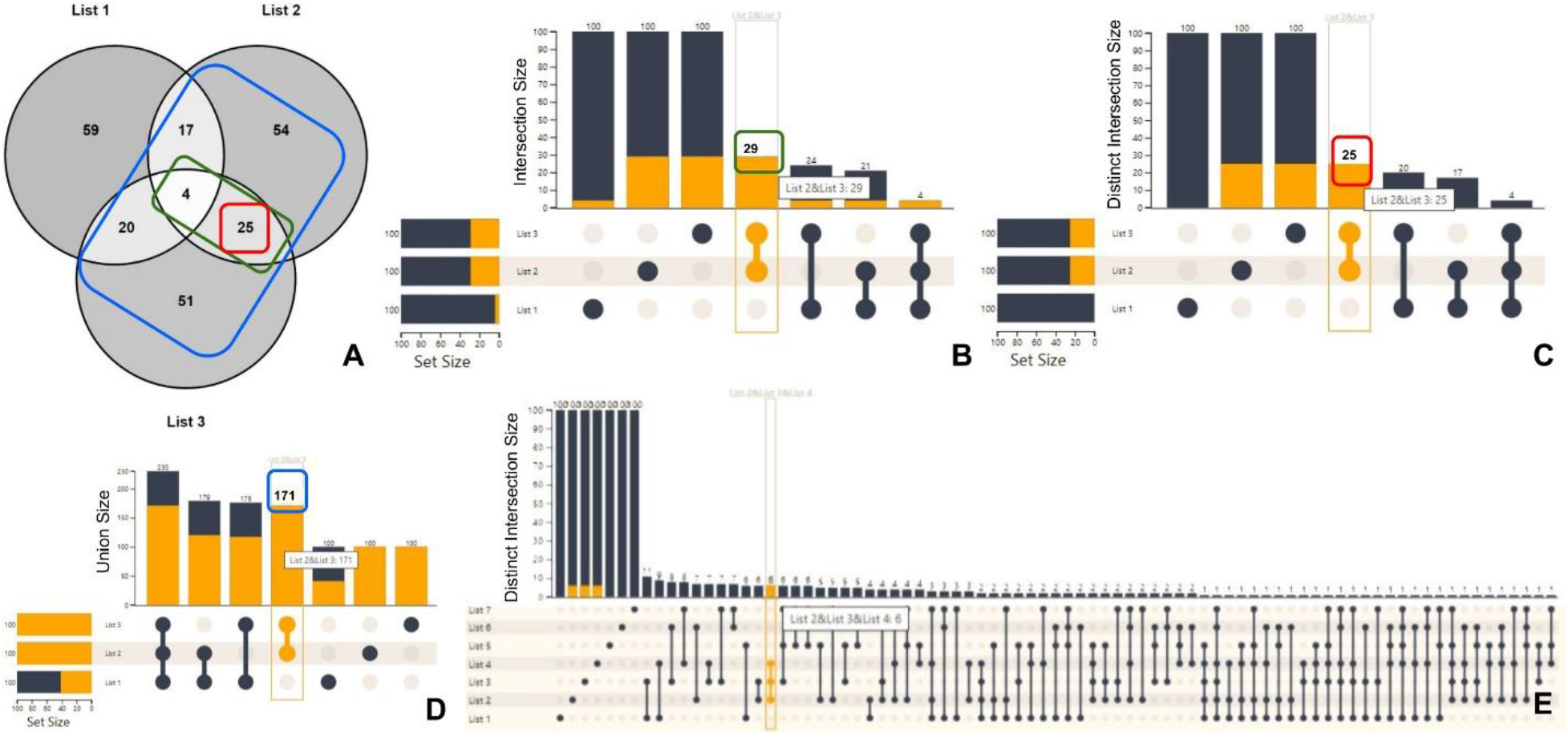
UpSet plot vs Venn diagrams. (A) Intersection of the three gene lists (100 genes each) shown in a Venn diagram. (B) The UpSet plot *Intersection* option visualizes the total number of common elements among the selected sets, even though they may also participate in other sets. For example, lists 2 and 3 contain 29 common genes (green rectangle), with 25 being shared only between them, and 4 also shared with List 1, as it can be seen in (A). (C) The *distinct intersections* option visualizes the common number of genes, among the chosen sets, which do not exist in any other set. This option is the closest to a Venn diagram. For example, lists 2 and 3 share 25 distinct genes (red rectangle). (D) The *union* option appends the unique elements among the chosen sets and creates all possible combinations. For example, the combination of lists 2 and 3 results in 171 total unique genes (blue rectangle). (E) A ‘*distinct intersections’* UpSet plot example with 7 lists, which cannot be visualized as a Venn diagram.

An UpSet plot consists of two axes and a connected-dot matrix. The vertical rectangles represent the number of elements participating in each list combination. The connected-dots matrix indicates which combination of lists corresponds to which vertical rectangle. Finally, the horizontal bars (Set Size) denote the participation of hovered objects (from the vertical rectangles) in the respective lists. In its current version, FLAME supports four UpSet plot modes: *i) intersections, ii) distinct intersections, iii*) *distinct elements per file* and *iii) unions*. The *intersections* mode creates all file combinations as long as they share at least one element, allowing an element to participate in more than one combination. The *distinct intersections* mode creates file combinations only for distinct elements that do not participate in other lists. The *distinct elements per file* mode shows the distinct elements of each input list. The *unions* mode constructs all available file combinations, showing their total unique elements. Notably, on mouse-hover over each UpSet plot element combination (vertical rectangles), FLAME presents the respective genes in a table, while on a mouse click the user can append the selected list of files with a new file containing the selected genes and process it separately.

### Functional enrichment

Once an input file or an UpSet plot column (*intersection*/*union* of sets) has been selected, FLAME takes advantage of the g:Profiler library (3) and aGOtool API (11) to offer functional enrichment analysis for a list of 197 organisms.

In detail, g:Profiler is used for the identification of enriched functional terms from Gene Ontology (19), pathways from KEGG (20), Reactome (22) and WikiPathways (23), protein complexes from CORUM (25), expression data from Human Protein Atlas (26), regulatory motifs from TRANSFAC (27) and miRTarBase (28), and phenotypes from the Human Phenotype Ontology (29). Similarly, aGOtool is used for the identification of enriched terms from the UniProt keyword classification system (30), protein families and domains from Pfam (31) and InterPro (32), as well as human diseases from the DISEASES database (33).

g:Profiler and aGOtool test for statistically significant enrichment to compare the user’s input gene list to a background set from organism-specific genes annotated in the Ensembl database (34) and UniProt Reference Proteomes (30), respectively. In the case of g:Profiler, the resulting *p*-values are corrected for multiple testing using g:SCS, Bonferroni correction or Benjamini-Hochberg false discovery rate (FDR), whereas in the case of aGOtool, *p*-values are corrected using Bonferroni correction or FDR. Notably, the reported lists with the enrichment results can be shrinked or expanded using the aforementioned parameters as thresholds.

Enrichment analysis is basically performed using ENSEMBL identifiers, while, based on the user’s choice, results can be reported as Entrez, UniProt (30), EMBL (35), ENSEMBL (34), ChEMBL (36), WikiGene (37) and RefSeq (38) identifiers. Notably, conversion to ENSEMBL identifiers is done internally at the backend, regardless of the identifier type imported by the user.

### Literature enrichment

In addition to functional enrichment, FLAME enables literature enrichment analysis for a selected gene list, via the aGOtool API. To this end, FLAME allows users to retrieve scientific articles which are tightly connected to the genes/proteins provided in the uploaded input files. The literature enrichment analysis concept is very similar to the one of the enrichment analysis and aims to aid users to identify scientific publications of relevance to a given gene/protein list.

The publication enrichment analysis is based on the aGOtool, which uses a text corpus of all PubMed abstracts and full-text open access articles from PubMed Central. These documents are processed by OnTheFly’s (39) or EXTRACT’s (40) underlying Named Entity Recognition (NER) tagger (41) on a weekly basis for the identification of biomedical entities (genes/proteins, chemical compounds, organisms, tissues, environments, diseases, phenotypes and Gene Ontology terms). As a result, all documents are being automatically annotated with the genes mentioned within them, thus like in functional enrichment, turning every document into a ‘gene set’.

### Protein-protein interaction analysis

FLAME offers the capability to construct and visualize interactive PPIs for a set of 197 organisms using the STRING API (24). Users may submit their gene list and visualize the results as networks, with the interacting entities being presented as nodes and their interactions as edges. In the online version, FLAME allows a maximum of 500 proteins per request, a limitation that can be bypassed by downloading the application from the GitHub repository and running it locally after editing the value of the corresponding variable (STRING_LIMIT configuration variable in global.R). We note that for each input gene, we only keep one converted Ensembl protein identifier.

STRING supports both physical or functional interactions. Physical interactions refer to proteins which are part of the same biomolecular complex, whereas functional interactions refer to proteins which are involved in the same pathway or biological process. Through FLAME, users can select whether to visualize the *full* set of interactions (both physical and functional), or just the *physical sub-network*. Users can also adjust the *interaction score* and apply a cutoff on the edges. In addition, users can choose between the *evidence* or the *confidence* mode. In the first case, a multi-edged graph is drawn, in which each edge shows the evidence channel (e.g. fusion event, co-expression and text mining), and the information comes from (42), whereas in the second case, the thickness of the edge reflects the interaction score. While the resulting network is presented in a separate Network Viewer panel, one can export a network as an image or as a tab-delimited file to be visualized by external viewers (43–45), or get redirected in STRING’s original source for further analysis.

### Visual analytics and interactive visualization

FLAME offers various interactive plots and visualization options for reporting results. Functional enrichment results produced by g:Profiler and aGOtool, as well as literature enrichment results produced by aGOtool can be shown as tables, scatter plots, barcharts, heatmaps and networks.

Resulting lists of functional terms are initially reported as interactive searchable tables displaying details about each functional term. One can expand each row of the table to see which of the identified genes/proteins were found to be associated with the functional term. For example, in the case of a KEGG pathway, one can see how many proteins or genes were found to be related to it and get redirected to the KEGG repository to see the actual schema of the pathway in a static form with all of the detected genes/proteins highlighted.

In the case of g:Profiler only, an ‘adjusted to the selected data sources’ interactive Manhattan plot is offered for a clearer overview. In this plot, functional terms are organized along the x-axis and colored by their data source, whereas the y-axis shows the significance (*p*-value) of each term. Hovering over a data point generates a popup window with key information about the functional term. By selecting a set of points using a lasso or a rectangle, the Manhattan plot will be redrawn showing information about the selected items only. Upon selection, the corresponding tables will be automatically updated.

In all of the enrichment-type analyses offered by FLAME, for every source, the most significant functional terms will be shown in a bar chart or a scatter plot, which the user can further customize to adjust the desired number of terms analyzed. Bar charts are sorted according to the enrichment score or the *p*-value. Finally, on mouse hovering, a tooltip with key information about the functional term will be shown.

Heatmap and network visualization options complement each other and come with two different execution modes. In the first case, genes are plotted against functional terms, while in the second case functional terms are plotted against themselves (square matrix). In the case of heatmaps, a color-scheme is used to capture the enrichment or the statistical score for a particular functional term. In the second mode (functional terms vs functional terms), a cell value depicts a similarity capturing the number of common genes. This is calculated as the summation of the unique common genes between a pair of functional terms divided by the number of total unique genes found to be associated with the functional terms. All of the heatmaps are fully interactive and one can zoom in and isolate an area of interest, swap the x and y axes and adjust the number of elements which will be shown. Notably, all heatmaps are clustered after applying a hierarchical clustering method and export options are also supported.

In addition to heatmaps, a three-mode network visualization is offered. In the first case, nodes represent genes and/or functional terms (colored differently), whereas edges represent the similarity scores between them, as explained before (weighted networks). In the second case, nodes represent the functional terms, while edges reflect the number of common genes between them. Finally, in the third mode, nodes represent genes which are connected based on the number of common functions or processes they are involved in.

While a heatmap is a preferable option for observing all possible pairwise similarities, in the second network mode, users can apply an edge-cutoff based on the similarity score to reduce the network’s density, and make visualization more appealing and more informative. Notably, networks are fully interactive and one can zoom in/out, adjust the layout accordingly or export the network in various file formats. Examples of all of the aforementioned visualizations are shown in Figure 2.

**Figure 2.**
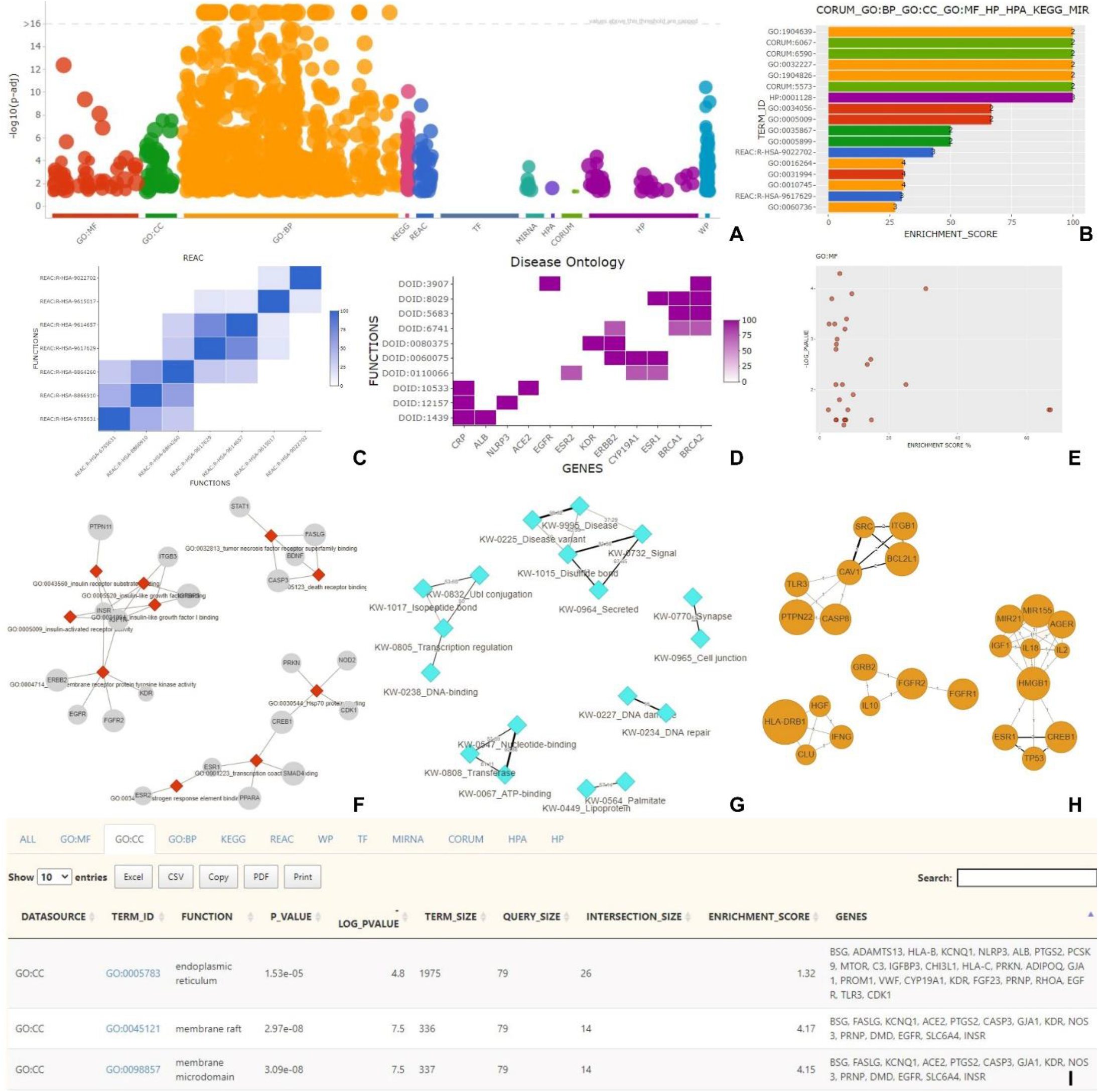
FLAME’s interactive visualization options. (A) g:Profiler’s result overview in a Manhattan plot. (B) A barchart, incorporating the top 17 results after combining different sources like GO Biological Process, GO Cellular Component, GO Molecular Function, phenotypes, protein complexes, pathways and miRNAs. (C) A hierarchically clustered heatmap with pairwise similarities between reactome terms. The color intensity reflects the common genes between each pair. (D) A heatmap showing genes (x-axis) associated with Disease Ontology terms (y-axis). (E) A scatter plot of enriched GO Molecular Function terms. The x-axis represents the enrichment score, while the y-axis the statistical significance. (F) An unweighted network consisting of GO Molecular Function functional terms and their related genes. (G) A weighted network of pairwise UniProt keywords returned by aGOtool. Edge weights are proportional to the number of common genes between each pair. (H) A gene-gene network based on common GOs, where an edge width is proportional to the number of common biological processes (BP) between two genes. (I) An interactive table with enrichment analysis results reported in different tabs (source of the active tab shown: Gene Ontology Cellular Component).

### Gene ID conversion and Orthology search

Frequently, different databases and tools accept varying gene/protein identifiers as their input. Thereby, the ID mapping creates a bottleneck for the non-expert end-users. To aid researchers in overcoming this problem, FLAME utilizes g:Profiler’s converters to allow i) cross-database ID conversions and ii) cross-species ID conversions (orthologs). In detail, FLAME allows ID conversions between well-known name-spaces, such as Entrez Gene, Uniprot, ChEMBL, ENSEMBL and RefSeq, as well as among the 197 different organisms supported by FLAME.

### Implementation

FLAME is mainly written in R and JavaScript. The R/Shiny package was used for the GUI implementation and the interoperability between R and Javascript. UpSet plots have been implemented with the use of R/upsetjs library. Functional enrichment analyses are offered by R/gprofiler2 library and aGOtool API. The latter is also used for literature enrichment analysis. Networks are stored as igraph objects (46) and visualized with the visNetwork library. Scatter and Manhattan plots are generated with the help of plotly library (47), bar plots with ggplot (48) and heatmaps via the heatmaply library (49). Interactive tables are generated through the DT library. Finally, network analysis is performed with the employment of STRING API.

### Integration with other applications

FLAME can be called from other applications with a simple get request. Gene names must be encoded in the URL (url_genes variable) and be comma-separated (,), whereas multiple gene lists must be separated by the semicolon symbol (;). Lists will appear as uploaded files. A simple URL example encoding three lists is: flame.pavlopouloslab.info/?url_genes=MCL1,TTR;APOE,ACE2;TLR4,HMOX1,TP73

## CASE STUDY

We have tested the capacity of FLAME for knowledge discovery by re-analyzing gene expression data associated with colitis-propelled carcinogenesis (CAC) in mice (50). In this model, tumorigenesis develops following a single application of the carcinogen azoxymethane (AOM) combined with four cycles of dextran sodium sulfate (DSS) administration that causes chronic colitis. However, inflammation, tissue damage, dysplasia and cancer are manifested in the distal but not the proximal part of the large intestine. To gain insight into the biological basis of this intriguing phenomenon, we have previously interrogated the transcriptome of proximal and distal colon, and reported intrinsic differences in gene expression which are augmented during CAC. These analyses also provided evidence that lipid metabolic pathways operating in the proximal part of the large intestine mediate resistance to experimental colitis and CAC (50).

Herein, we have focused on exploring gene ontologies and pathways that may mediate sensitivity to CAC. We reasoned that transcripts that are upregulated at both the “early” (i.e. 2 DSS cycles) and “late” (i.e. 4 DSS cycles) stages of AOM/DSS-induced carcinogenesis in the distal part of the colon and are not detected in the proximal region must play a prominent role in CAC. We have identified 165 transcripts belonging to this intersection by using the “UpSet Plot” option of FLAME (Figure 3A and Suppl. Table 1) and termed this gene set ‘the susceptibility-associated gene signature’, SAS.

**FIGURE 3:**
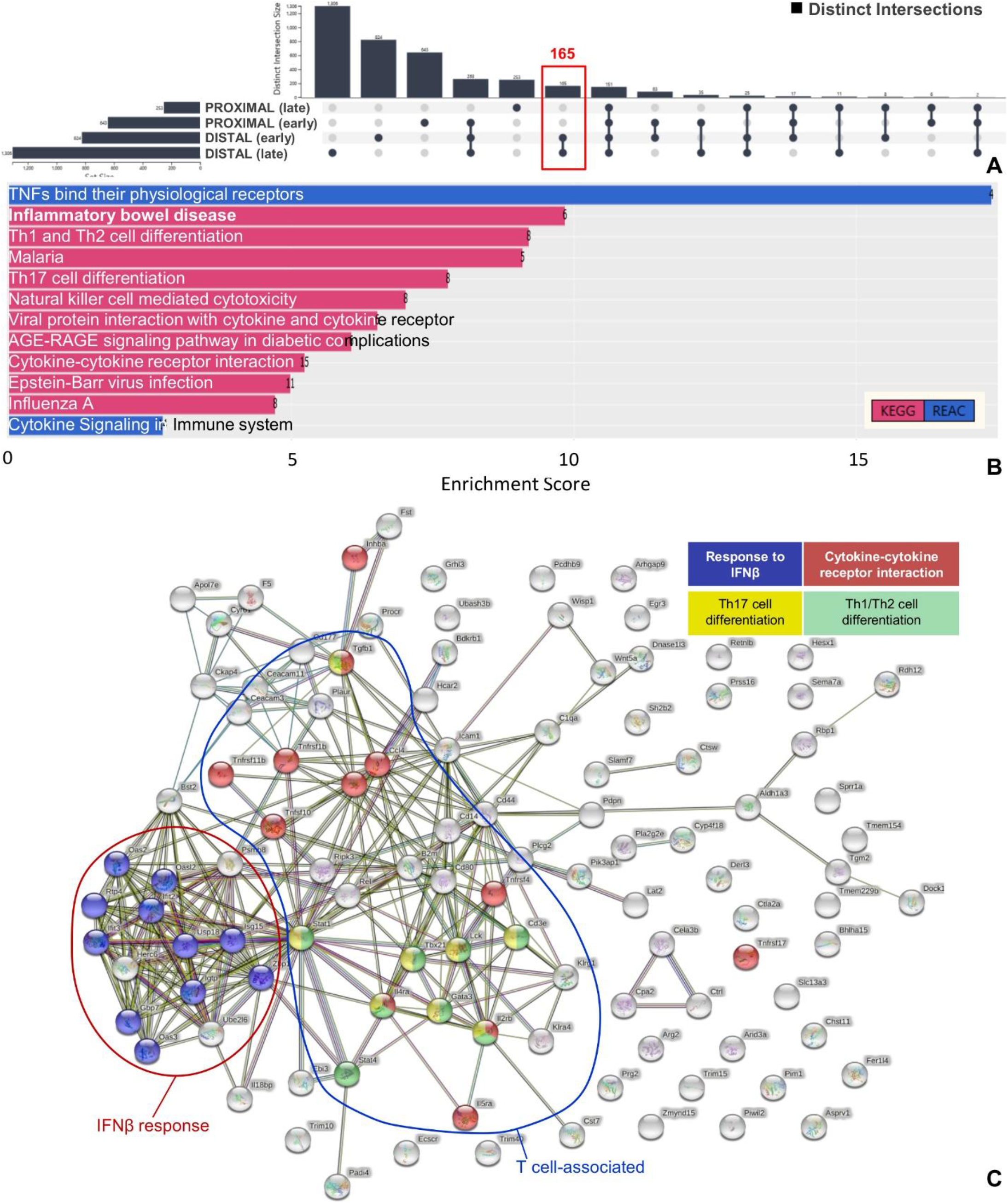
(A) UpSet plot of distinct intersections of transcripts upregulated at the “early” (i.e. 2 DSS cycles) and “late” (i.e. 4 DSS cycles) stages of AOM/DSS-induced carcinogenesis in the disease-resistant proximal and disease-sensitive distal part of the colon. The “susceptibility-associated gene signature’, SAS, comprising 165 transcripts was used for gene ontology and pathway enrichment. (B) KEGG and Reactome pathway enrichment of SAS transcripts. (C) Analysis of putative protein-protein interactions through the STRING option of FLAME. A cluster of interacting components of interferon beta (IFNβ) response and of T cell-associated immunity, including Th1/Th2 differentiation, Th17 differentiation and cytokine-cytokine receptor interaction, each represented by a different color, is shown.

To interpret SAS transcripts as biological functions and pathways, we combined KEGG and REACTOME pathway enrichment analysis using FLAME. Reassuringly, we found “Inflammatory Bowel Disease” to be among the most significantly enriched pathways (Figure 3B and Suppl. Table 2). Other pathways found to be significantly enriched in SAS were related to immune function, including T helper 1 (Th1)/Th2 differentiation, Th17 cell differentiation, Natural Killer (NK) cell-mediated cytotoxicity and cytokine/chemokine signaling which align with the pivotal role of unresolved inflammation in promoting intestinal cancer (51). Indeed, Th17 lymphocytes give rise to pathogenic Th1 cells implicated in colitis (52) and when activated by TGFβ1 (a SAS gene set cytokine; Suppl. Table 1), Th17 cells promote CAC (53). Moreover, chronic experimental colitis is associated with dual Th1 and Th2 cytokine profile (54) and there is evidence to suggest that Th2-driven colonic inflammation enhances the formation of colorectal tumors (55). We also note the significant enrichment of SAS for the pro-inflammatory TNF pathway which represents a major therapeutic target for inflammatory bowel diseases.

Gene ontology (GO) enrichment was also performed on the FLAME platform. SAS was found to be enriched for biological processes that were mostly related to T cell-mediated immunity and cytokine synthesis (Suppl. Figure 1A and Suppl. Table 3). The GO enrichment for molecular functions (Suppl. Figure 1B and Suppl. Table 3) identified several terms related to *cytokine and chemokine activity*, including TNF, and to *death receptor signaling* which has been implicated in CAC (56). A gene-gene network was also constructed using FLAME to uncover clusters of SAS genes with common molecular functions (Suppl. Figure 1C). Analysis of putative protein-protein interactions (physical and functional associations) through the STRING option of FLAME uncovered a cluster of interacting components of T cell-associated immunity and a cluster of interferon beta (IFNβ) response proteins implicated in epithelial regeneration upon DSS-induced tissue damage (57) (Figure 3C).

To gain putative mechanistic insights into the aforementioned biological functions and pathway data, we used the FLAME platform to perform transcription factor motif enrichment and analyze transcription factor networks versus SAS transcripts. The results showed a cluster of interferon regulatory factor (IRF) family members (Suppl. Figure 1D and Suppl. Table 4), which may be linked to both the enrichment of SAS in IFNβ components and T helper cell differentiation. Another transcription factor cluster was associated with SMADs which have been implicated in CAC, in part downstream of TGFβ1 (58, 59). Overall, the aforementioned analyses underscore the practical utility of FLAME to rapidly process, analyze and visualize gene expression data and hence assist knowledge discovery.

## DISCUSSION

FLAME is a web application targeting to effectively complement existing enrichment tools and make gene list analysis, information extraction, knowledge integration and visualization much easier, more comprehensive, accurate and reliable. It combines functionalities from g:Profiler, aGOtool and STRING to cover a wide spectrum of analyses, such as functional, literature enrichment and network analysis. FLAME is designed for non-experts and via the engagement of UpSet plots, it allows users to manipulate, compare and handle multiple gene lists prior to any type of analysis. Moreover, it follows a visual analytics approach, thus offering users various options for visualizing their results and adjusting the corresponding views upon parameterization to generate publication-accepted figures. In a future version, we expect to expand FLAME’s capabilities by supporting more organisms and integrating a greater plethora of sources.

## CONCLUSION

FLAME is a user-friendly visual analytics web tool specifically designed for combining gene lists prior to enrichment analysis and creating informative interactive visualizations for the publication-required presentation of obtained results. It covers a wide spectrum of analyses and comes with an easy-to-use interface, aiming to successfully complement and cover gaps of existing state-of-the-art enrichment tools, and reach out to many users varying from non-experts to highly specialized bioinformaticians.

## AUTHOR CONTRIBUTIONS

F.T. and E.K. wrote parts of the manuscript and implemented most of the tool. F.B. implemented parts of the tool and the manuscript, and helped in the debugging process. D.J.S. and A.G.E. tested the tool thoroughly and provided valuable feedback. A.G.E. provided the case study. G.A.P. conceptualized and supervised the whole project, and provided feedback throughout its development.

## AVAILABILITY

FLAME is available at: http://flame.pavlopouloslab.info. The source code and instructions about the necessary dependencies can be found at: https://github.com/PavlopoulosLab/FLAME.

## ACKNOWLEDGEMENTS AND FUNDING

This article was supported by the matching funds from the grant supported by the European Union’s Horizon 2020 research and innovation programme under the Marie Skłodowska-Curie grant agreement No 838018. The study was also supported by the Hellenic Foundation for Research and Innovation (H.F.R.I.) under the “First Call for H.F.R.I. Research Projects to support faculty members and researchers and the procurement of high-cost research equipment grant”, Grant ID: 1855-BOLOGNA. We also acknowledge support of this work by the project “The Greek Research Infrastructure for Personalised Medicine (pMedGR)” (MIS 5002802), which is implemented under the Action “Reinforcement of the Research and Innovation Infrastructure”, funded by the Operational Programme “Competitiveness, Entrepreneurship and Innovation” (NSRF 2014-2020) and co-financed by Greece and the European Union (European Regional Development Fund). Finally, the study was financially supported by the Operational Program Competitiveness, Entrepreneurship and Innovation, NSRF 2014-2020, Action code: MIS 5002562, co-financed by Greece and the European Union (European Regional Development Fund).

---

**Supplementary Figure 1:**
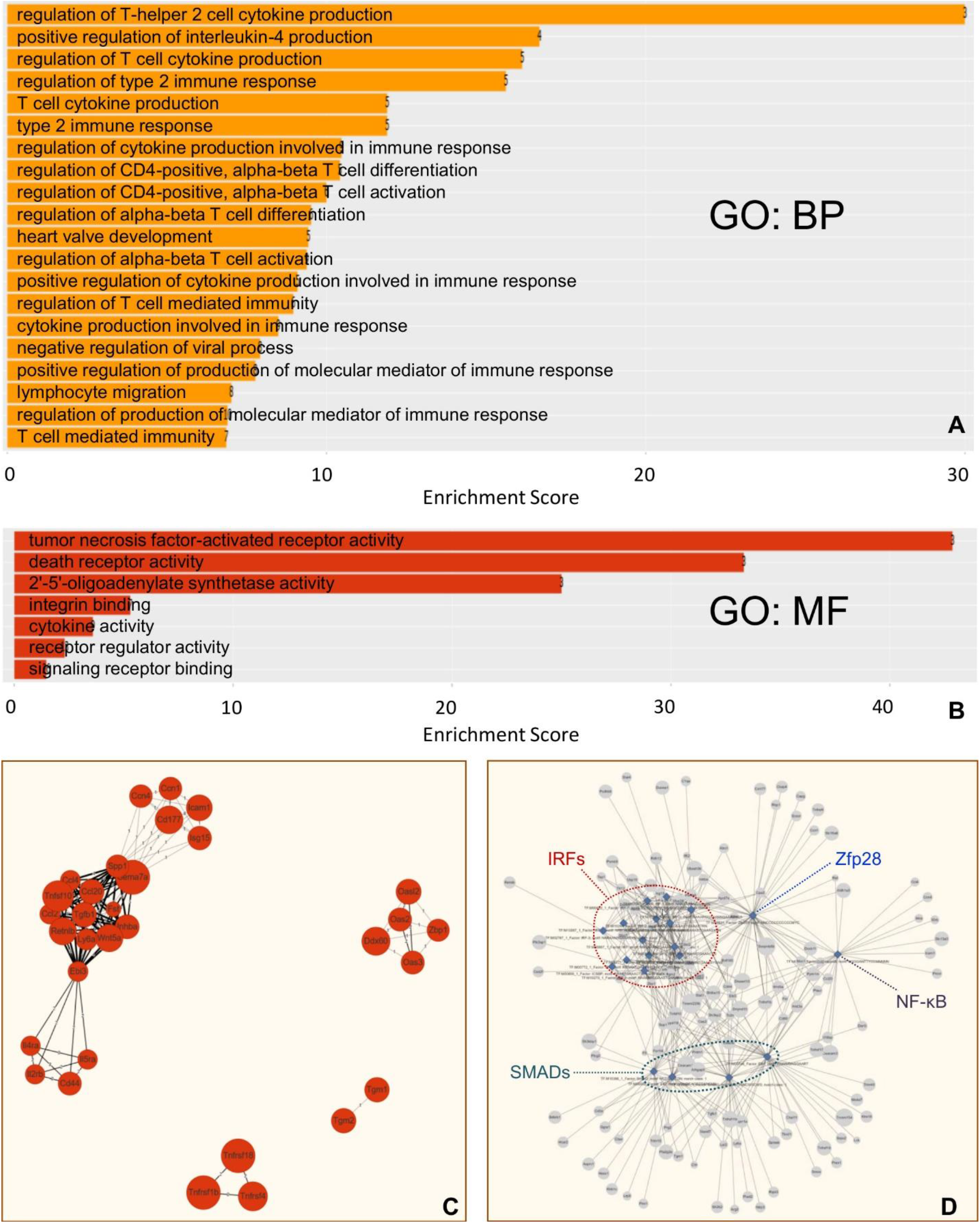
(A) Gene Ontology (GO) enriched biological processes (BP) of “susceptibility-associated gene signature”, SAS. (B) GO enriched molecular functions (MF) of SAS. (C) A MF-enriched gene-gene network, where an edge width is proportional to the number of common molecular functions between two genes. (D) An unweighted network of transcription factors (Functions vs Genes) predicted to be involved in the regulation of SAS genes.

**Supplementary Table 1.**
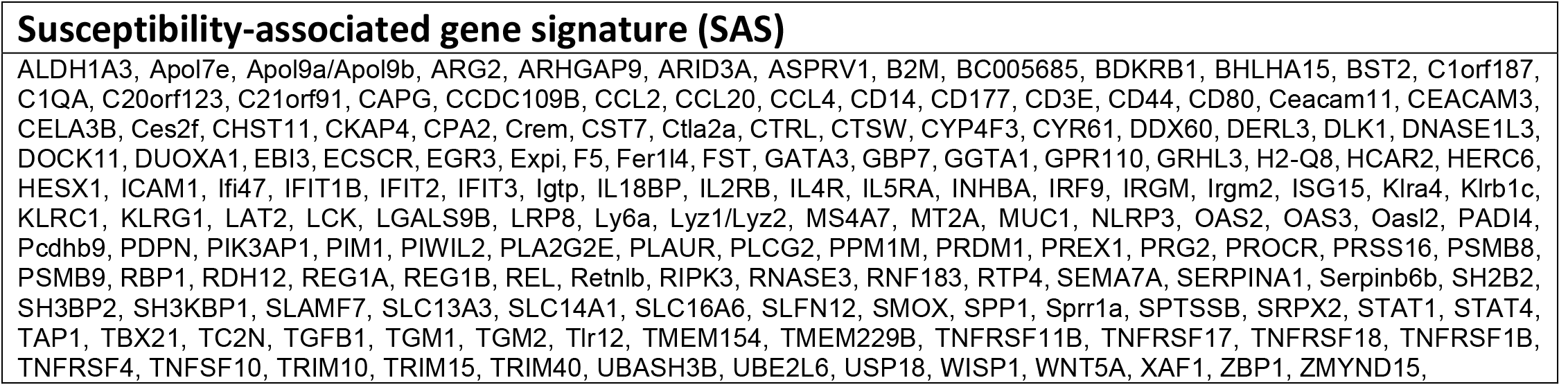

**Supplementary Table 2.**

| KEGG/REACTOME PATHWAYS |  |  |  |  |  |  |  |  |
| --- | --- | --- | --- | --- | --- | --- | --- | --- |
| DATASOURCE | TERM_ID | FUNCTION | P_VALUE | OG_PVALUE | TERM_SIZE | QUERY_SIZE | INTERSECTION_SIZE | ENRICHMENT_SCORE |
| REAC | REAC:R-MMU-5669034 | TNFs bind their physiological receptors | 7.75e-03 | 2,1 | 23 | 69 | 4 | 17,39 |
| KEGG | KEGG:05321 | Inflammatory bowel disease | 2.53e-03 | 2,6 | 61 | 85 | 6 | 9,84 |
| KEGG | KEGG:04658 | Th1 and Th2 cell differentiation | 1.87e-04 | 3,7 | 87 | 85 | 8 | 9,2 |
| KEGG | KEGG:05144 | Malaria | 1.76e-02 | 1,8 | 55 | 85 | 5 | 9,09 |
| KEGG | KEGG:04659 | Th17 cell differentiation | 6.68e-04 | 3,2 | 103 | 85 | 8 | 7,77 |
| KEGG | KEGG:04650 | Natural killer cell mediated cytotoxicity | 1.41e-03 | 2,8 | 114 | 85 | 8 | 7,02 |
| KEGG | KEGG:04061 | Viral protein interaction with cytokine and cytokine receptor | 2.53e-02 | 1,6 | 92 | 85 | 6 | 6,52 |
| KEGG | KEGG:04933 | AGE-RAGE signaling pathway in diabetic complications | 3.77e-02 | 1,4 | 99 | 85 | 6 | 6,06 |
| KEGG | KEGG:04060 | Cytokine-cytokine receptor interaction | 1.08e-05 | 5 | 287 | 85 | 15 | 5,23 |
| KEGG | KEGG:05169 | Epstein-Barr virus infection | 9.61e-04 | 3 | 221 | 85 | 11 | 4,98 |
| KEGG | KEGG:05164 | Influenza A | 2.39e-02 | 1,6 | 170 | 85 | 8 | 4,71 |
| REAC | REAC:R-MMU-1280215 | Cytokine Signaling in Immune system | 1.13e-02 | 1,9 | 515 | 69 | 14 | 2,72 |

**Supplementary Table 3.**

| SAS ENRICHMENT FOR GENE ONTOLOGY BIOLOGICAL FUNCTIONS |  |  |  |  |  |  |  |  |
| --- | --- | --- | --- | --- | --- | --- | --- | --- |
| DATASOURCE | TERM_ID | FUNCTION | P_VALUE | -LOG_PVALUE | TERM_SIZE | QUERY_SIZE | INTERSECTION_SIZE | ENRICHMENT_SCORE |
| GO:BP | GO:2000551 | regulation of T-helper 2 cell cytokine production | 4.89e-02 | 1,3 | 10 | 136 | 3 | 30 |
| GO:BP | GO:0032753 | positive regulation of interleukin-4 production | 2.52e-02 | 1,6 | 24 | 136 | 4 | 16,67 |
| GO:BP | GO:0002724 | regulation of T cell cytokine production | 2.41e-03 | 2,6 | 31 | 136 | 5 | 16,13 |
| GO:BP | GO:0002828 | regulation of type 2 immune response | 2.84e-03 | 2,5 | 32 | 136 | 5 | 15,62 |
| GO:BP | GO:0002369 | T cell cytokine production | 1.14e-02 | 1,9 | 42 | 136 | 5 | 11,9 |
| GO:BP | GO:0042092 | type 2 immune response | 1.14e-02 | 1,9 | 42 | 136 | 5 | 11,9 |
| GO:BP | GO:0002718 | regulation of cytokine production involved in immune response | 6.68e-06 | 5,2 | 86 | 136 | 9 | 10,47 |
| GO:BP | GO:0043370 | regulation of CD4-positive, alpha-beta T cell differentiation | 2.23e-02 | 1,7 | 48 | 136 | 5 | 10,42 |
| GO:BP | GO:2000514 | regulation of CD4-positive, alpha-beta T cell activation | 3.76e-03 | 2,4 | 60 | 136 | 6 | 10 |
| GO:BP | GO:0046637 | regulation of alpha-beta T cell differentiation | 5.02e-03 | 2,3 | 63 | 136 | 6 | 9,52 |
| GO:BP | GO:0003170 | heart valve development | 3.64e-02 | 1,4 | 53 | 136 | 5 | 9,43 |
| GO:BP | GO:0046634 | regulation of alpha-beta T cell activation | 1.79e-05 | 4,7 | 96 | 136 | 9 | 9,38 |
| GO:BP | GO:0002720 | positive regulation of cytokine production involved in immune response | 4.37e-02 | 1,4 | 55 | 136 | 5 | 9,09 |
| GO:BP | GO:0002709 | regulation of T cell mediated immunity | 7.22e-03 | 2,1 | 67 | 136 | 6 | 8,96 |
| GO:BP | GO:0002367 | cytokine production involved in immune response | 4.29e-05 | 4,4 | 106 | 136 | 9 | 8,49 |
| GO:BP | GO:0048525 | negative regulation of viral process | 4.51e-04 | 3,3 | 101 | 136 | 8 | 7,92 |
| GO:BP | GO:0002702 | positive regulation of production of molecular mediator of immune response | 5.24e-04 | 3,3 | 103 | 136 | 8 | 7,77 |
| GO:BP | GO:0072676 | lymphocyte migration | 1.14e-03 | 2,9 | 114 | 136 | 8 | 7,02 |
| GO:BP | GO:0002700 | regulation of production of molecular mediator of immune response | 5.21e-05 | 4,3 | 145 | 136 | 10 | 6,9 |
| GO:BP | GO:0002456 | T cell mediated immunity | 6.77e-03 | 2,2 | 102 | 136 | 7 | 6,86 |
| SAS ENRICHMENT FOR GENE ONTOLOGY MOLECULAR FUNCTIONS |  |  |  |  |  |  |  |  |
| DATASOURCE | TERM_ID | FUNCTION | P_VALUE | -LOG_PVALUE | TERM_SIZE | QUERY_SIZE | INTERSECTION_SIZE | ENRICHMENT_SCORE |
| GO:MF | GO:0005031 | tumor necrosis factor-activated receptor activity | 4.21e-03 | 2,4 | 7 | 136 | 3 | 42,86 |
| GO:MF | GO:0005035 | death receptor activity | 1.00e-02 | 2 | 9 | 136 | 3 | 33,33 |
| GO:MF | GO:0001730 | 2'-5'-oligoadenylate synthetase activity | 2.58e-02 | 1,6 | 12 | 136 | 3 | 25 |
| GO:MF | GO:0005178 | integrin binding | 1.12e-02 | 2 | 132 | 136 | 7 | 5,3 |
| GO:MF | GO:0005125 | cytokine activity | 1.75e-02 | 1,8 | 251 | 136 | 9 | 3,59 |
| GO:MF | GO:0030545 | receptor regulator activity | 3.50e-02 | 1,5 | 564 | 136 | 13 | 2,3 |
| GO:MF | GO:0005102 | signaling receptor binding | 2.95e-02 | 1,5 | 1780 | 136 | 26 | 1,46 |

**Supplementary Table 4.**
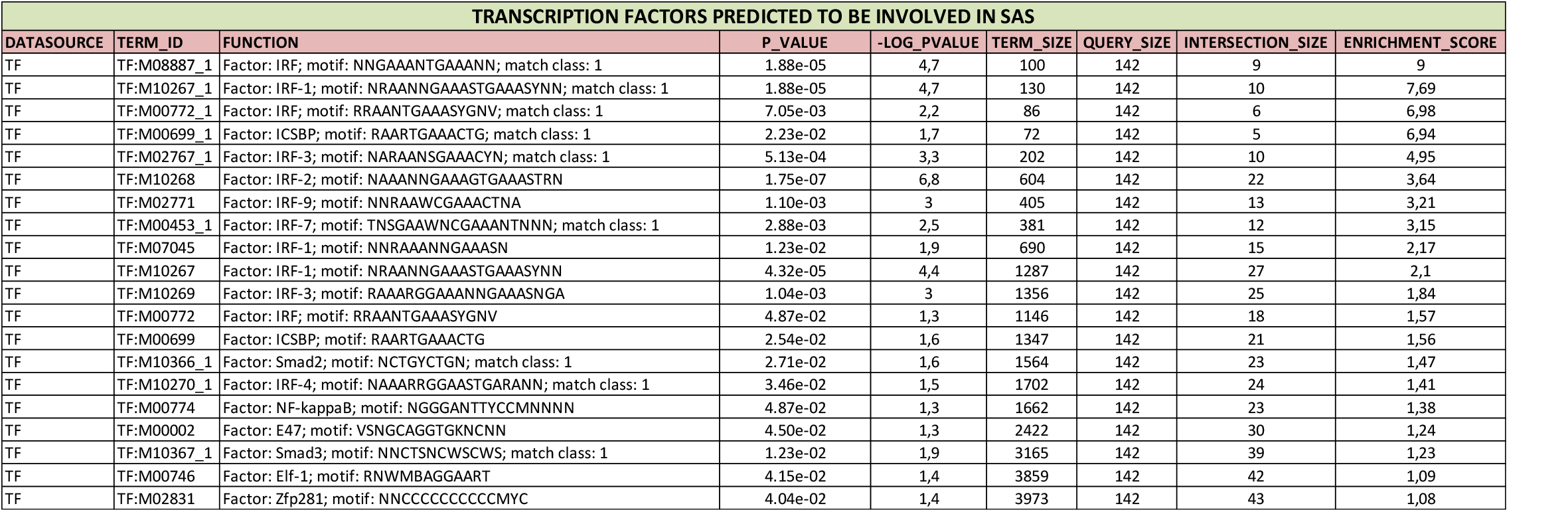

